# Membrane lipid reshaping underlies oxidative stress sensing by the mitochondrial proteins UCP1 and ANT1

**DOI:** 10.1101/2022.07.06.498870

**Authors:** Olga Jovanović, Ksenia Chekashkina, Sanja Škulj, Kristina Žuna, Mario Vazdar, Pavel V. Bashkirov, Elena E. Pohl

## Abstract

Oxidative stress and ROS are important players in the pathogenesis of several diseases. Besides the direct modification of proteins, ROS modify lipids with negative spontaneous curvature, such as phosphatidylethanolamine (PE), producing PE adducts and lysolipids. The formation of PE-adducts potentiates the protonophoric activity of the uncoupling protein 1 (UCP1), but the molecular mechanism remains obscure. Here, we connected the ROS-mediated lipid shape alteration with the membrane mechanical properties and function of UCP1 and adenine nucleotide translocase 1 (ANT1). We found that lysophosphatidylcholines (OPC and MPC) and PE adducts decrease a bending modulus in lipid bilayers and increase the protonophoric activity of both proteins. Furthermore, MD simulations revealed that modified PEs and lysolipids alter the membrane lateral pressure profile in the same direction and range, indicating that modified PEs act as lipids with positive spontaneous curvature. Both results indicate that oxidative stress decreases stored curvature elastic stress (SCES) in the lipid bilayer membrane. In conclusion, we demonstrate that UCP1 and ANT1 sense SCES and propose a new regulatory mechanism for the function of these proteins.

## Introduction

Lipid heterogeneity, observed between the membranes of cells and cell organelles, as well as between membrane leaflets and subdomains, is essential for proper cell functioning ^1, 2, 3, 4, 5^. The most abundant lipids in cell membranes are phospholipids, which differ in shape and spontaneous curvature (SC). Phosphatidylglycerol and phosphatidylcholine (PC), which spontaneously assemble into lamellar bilayer structures at physiological concentration of ions in water, have a cylindrical shape and negligible SC (≈0). Non-bilayer lipids or conical lipids such as phosphatidylethanolamine (PE; SC < 0) and lysolipids (lysoPC; SC > 0) form highly curved lipid structures such as HII phase (PE) or micelles. PE accounts for approximately 25% of all membrane lipids that significantly affect the mechanical properties of membranes ^6, 7^.

The embedding of conical lipids into a flat monolayer modifies the lateral pressure profile (LPP) so that the bending moment in the monolayer is no longer zero ^8^. Maintenance of the monolayer flatness leads to the induction of stored elastic curvature stress (SCES) ^9^. SCES induced by PE lipids supports the docking and insertion of membrane-associated/peripheral proteins ^7, 10, 11^, membrane budding ^12^, and membrane remodeling ^13, 14^. SCES is also suggested to influence the functioning of transmembrane proteins. However, only few studies have shown this relationship ^9, 15, 16, 17, 18, 19, 20^.

The role of conical lipids is especially apparent in the inner mitochondrial membrane (IMM), which contains the highest percentage of phospholipids with a negative SC (38%–45% PE and 15%–25% cardiolipin [CL]) ^21, 22^. Their shape supports the characteristic architecture of the IMM, which is necessary for the proper functioning of the mitochondrion ^23, 24, 25, 26^. Because PE and CL sense membrane curvature, curvature-driven redistribution of PE leads to its accumulation in strongly concave regions ^27, 28, 29, 30^. As an example, CL and PE support adenosine 5-triphosphate (ATP) synthase dimer formation, generating strong membrane curvature on the top of the cristae and thereby potentiating protein activity ^31, 32^.

Under oxidative stress conditions, membrane lipids undergo shape transformation. The products of reactive oxygen species (ROS), which comprise the biologically important RAs 4-hydroxy-2-nonenal (HNE) and 4-oxo-2-nonenal, covalently modify the PE head group and form RA-PE adducts ^33, 34, 35, 36^. RA-PE adducts increase the protonophoric activity of mitochondrial uncoupling protein 1 (UCP1), an essential player in non-shivering thermogenesis ^33^. Based on results obtained by mass spectrometry and molecular dynamic (MD) simulations ^37^, we previously suggested that the formation of RA-PE adducts reshapes PE, thereby changing the intrinsic curvature of this lipid. Another contribution to the transformation of the lipid shape during oxidative stress in mitochondria comes from the amplified action of membrane-bound phospholipase A2 (mPLA2) ^38, 39^, which converts membrane lipids to lysolipids (SC > 0) and free fatty acids (FFAs). The released FFAs represent raw material for the production of more RAs, which then target the headgroup of the PE and form more RA-PE adducts. As a result of these events, the ratio of lipids with opposite shapes in IMM can rapidly alter, leading to a modification of the LPP and the bending rigidity of the IMM ^40^. We hypothesized that the SCES shift caused by lipid shape transformation is a critical event by which IMM proteins (particularly uncoupling proteins) rapidly respond to the changes caused by oxidative stress.

To investigate the impact of ROS-modified lipids on mitochondrial protein function, we combined experimental measurements of membrane elastic properties and the total conductance (***G***_*m*_) of membranes containing recombinant UCP1 and ANT1 with calculations of membrane LPPs using MD simulations.

## Results

### 1. Impact of DOPE on membrane elastic parameters

First, we examined how the presence of PE influences the elastic properties of lipid bilayer membranes. To do so, we measured the radii (*r*_*NT*_) of nanotubes (NTs) pulled from a reservoir black lipid membrane (BLM) at different voltage biases (*U*) applied to the NT interior (Fig. 1A and Methods). The *r*_*NT*_ was extracted from a hyperbolic approximation of the measured conductance dependency on the change in NT length (Fig. 1B) at a fixed voltage.

**Figure 1.**
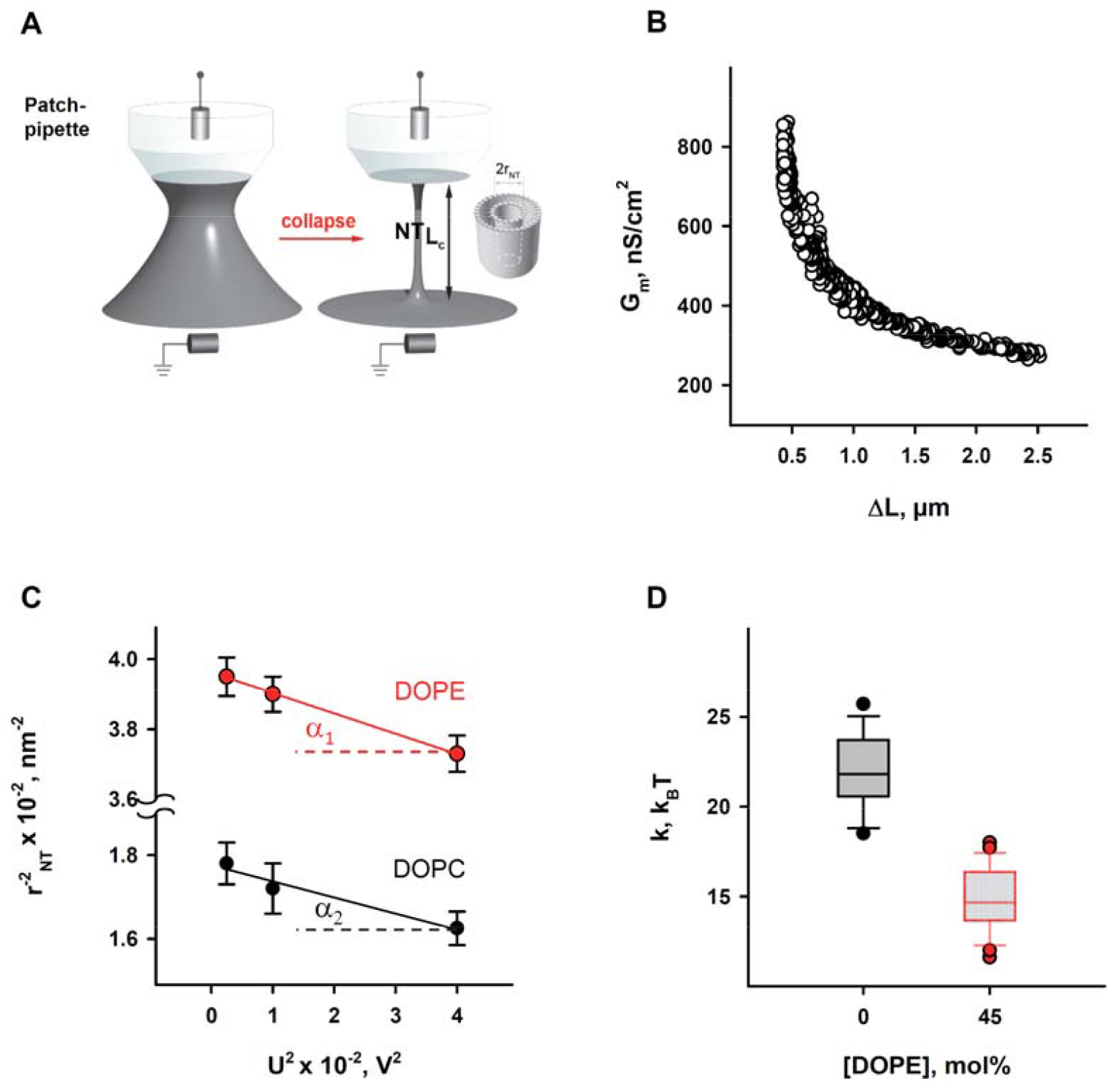
Determination of the elastic parameters of the lipid bilayer membrane based on nanotube (NT) pulling. A. Scheme of NTs pulled from a black lipid membrane (grey) and held by a patch-pipette (white). A voltage applied to the ends of the NTs induces an ion current (I_NT_) flowing through the NT interior. B. A representative measurement shows the dependence of membrane conductance (G_m_) (black circles) on the NT length change (ΔL), required for calculation of the NT radius (r_NT_) (see Methods). C. Dependence of the NT radius (r_NT_) on the applied voltage (U), obtained for the membrane lipid compositions DOPC:CL 90:10 (black) and DOPC:DOPE:CL 45:45:10 (red). The r_NT_ was estimated at different values of U applied to the ends of the NTs. Slopes (a) of the related linear regression provide the bending modulus k, while the lateral tension σ is estimated from the intercept with the ordinate axis. D. Box plot and distribution of the bending modulus k depending on the molar concentration of the phosphatidylethanolamine (DOPE). The buffer solution contained: 100 mM KCl, 10 mM HEPES, and 1 mM EDTA at pH 7.0 and T = 300 K. Data points represent the mean and standard deviation from more than 10 independent experiments.

To evaluate the impact of PE on membrane elastic parameters, we compared bilayer membranes comprising either (i) DOPC:DOPE:CL (45:45:10 mol%), mimicking the IMM, or (ii) DOPC:CL (90:10 mol%), as a model of a PE-free membrane. The r_NT_ was measured for both lipid compositions at U values varying from 50 to 200 mV. The membrane bending rigidity modulus (*k*) and lateral tension (*σ*) of BLMs were calculated from the linear regression of 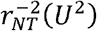 according to Eq. 1:

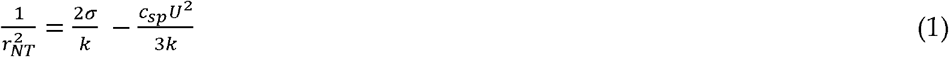

where c_sp_ stands for the specific membrane capacity.

We found that membranes containing 45 mol% DOPE were considerably less resistant to bending than DOPE-free membranes (Fig. 1D). The measured bending modulus decreased from *k*=22.1 ± 2.0 k_B_T for PE-free membranes to *k*=14.8 ± 1.8 k_B_T for PE-containing membranes.

This observation qualitatively confirmed the previously reported effect of membrane softening by PE ^41^, demonstrating that the addition of 30 mol% DOPE significantly reduced the *k*. On a molecular level, the effect was attributed to the curvature-driven redistribution of DOPE between NT monolayers and the reservoir membrane (BLM) during the formation of the NT. Conical lipids with a pronounced SC<0 tend to accumulate in the inner leaflet and to be depleted from the outer monolayer of the highly curved NT membrane, which remained connected to a flat reservoir membrane during the measurement ^29, 42^. Curvature-driven lipid redistribution resulted in a reduction in the apparent bending rigidity *k*, as given in Eq. 2 ^43^:

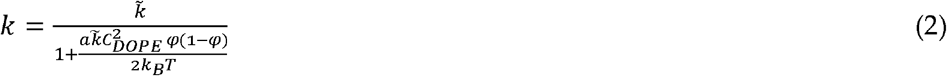

where 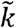 is the bending modulus of membranes with a fixed lipid composition, *a* – area per lipid (about 0.7 nm^2^), *C*_*DOPE*_ – spontaneous curvature of DOPE, and *ϕ* - the molar fraction of DOPE in the lipid reservoir.

We neglected the contribution of the CL shape to the apparent bending rigidity *k* because of its low amount (10 mol% of total lipid composition) and smaller SC (*C*_*CL*_ ≈ - 0.15 nm^-1^) ^44^. Thus, for PE-containing membrane (i), we considered 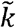 equal to the bending modulus measured for PE-free membranes and characteristic of DOPC membranes ^45^. Using Eq. (2), we calculated *C*_*DOPE*_ ≈ - 0.45 nm^-1^, which is in agreement with previously published data obtained for DOPE in the inverted HII phase ^46^, confirming the reduction in apparent bending rigidity due to curvature-composition coupling ^40, 47^. Notably, the lateral tension of membranes in contact with a reservoir that fixes the chemical potentials of compounds increases proportionally with the amount of DOPE following SCES induced by the packaging of conical lipids into a flat membrane. Indeed, we found the lateral tension of the membranes containing 50 mol% DOPE to be almost twice that of DOPE-free membranes (Fig. 3B), confirming the direct contribution of SCES to the lateral tension of membranes connected to a lipid reservoir ^40^.

### 2. Reactive aldehydes modify the elastic properties of the PE-containing lipid bilayer membranes

Next, we studied the effects of the modification of the lipids by reactive aldehydes (RAs) on the elastic properties of membranes. We incubated the reservoir membranes comprising different lipids, as specified in (i) and (ii), with a RA (4-hydroxy-2-hexenal [HHE], HNE, or ONE) at the concentrations of 0.5 - 0.7 mM. We found that HNE and ONE induced a pronounced reduction in the apparent bending modulus *k* in DOPE-containing membranes, resulting in *k*_*HNE*_ = 11.7 k_B_T for HNE and *k*_*ONE*_ = 7.1 k_B_T for ONE (Fig. 2A and Suppl. Fig. 1A). HHE had no detectable impact on *k* (Fig. 2A and Suppl. Fig. 1A). We explained the lack of the effect of HHE by its weaker adsorption to the lipid membrane ^33^. Remarkably, in the PE-free membranes, none of the RAs caused any measurable change in the bending modulus *k* (Fig. 2B and Suppl. Fig. 1B).

**Figure 2.**
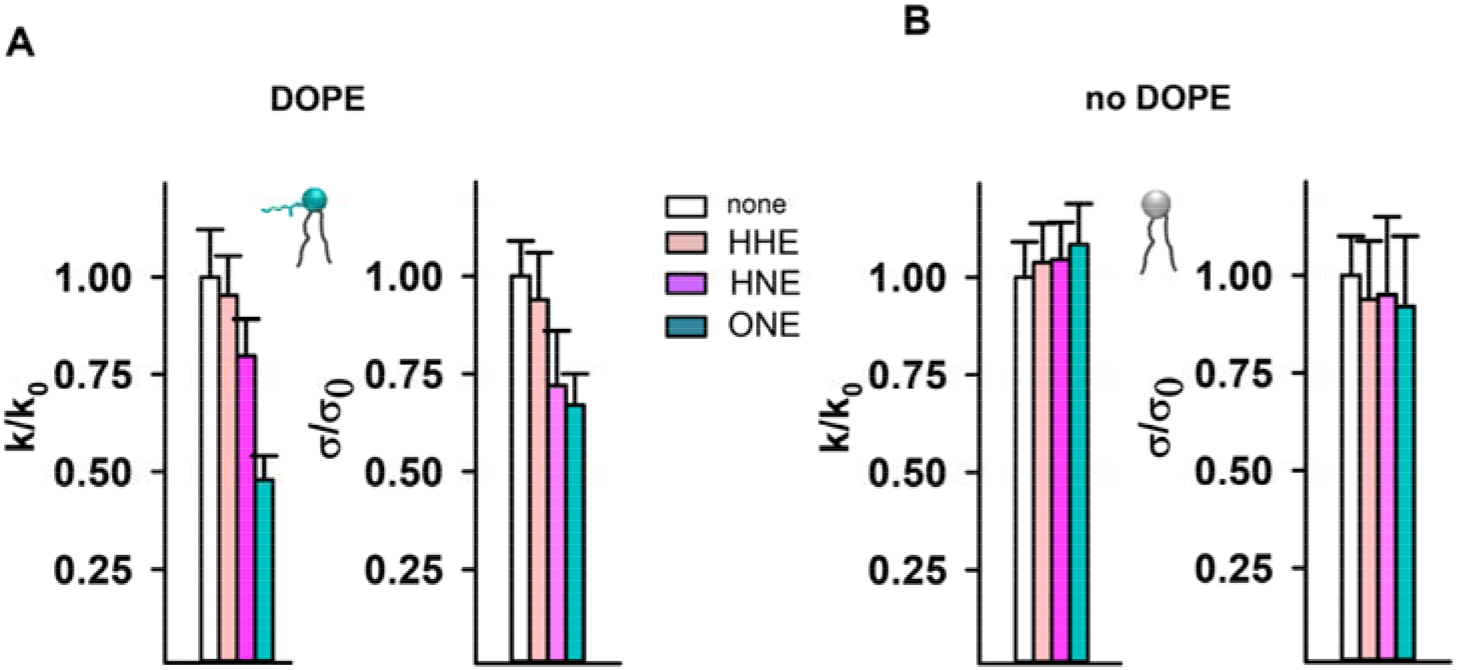
Elastic properties of the lipid bilayer membrane in the presence of reactive aldehydes (RAs). A. and B. Relative bending rigidity k/k_0_ and relative lateral tension σ/σ_0_ for lipid membranes composed with (A) and without (B) DOPE, incubated with the RAs 4-hydroxy-2-hexenal (HHE), 4-hydroxy-2-nonenal (HNE), and 4-oxo-2-nonenal (ONE). Membrane composition was as described in Figure 1. RAs were added in a concentration range 500 – 700 µM. Data points represent mean and standard deviation from more than 10 independent experiments.

**Figure 3.**
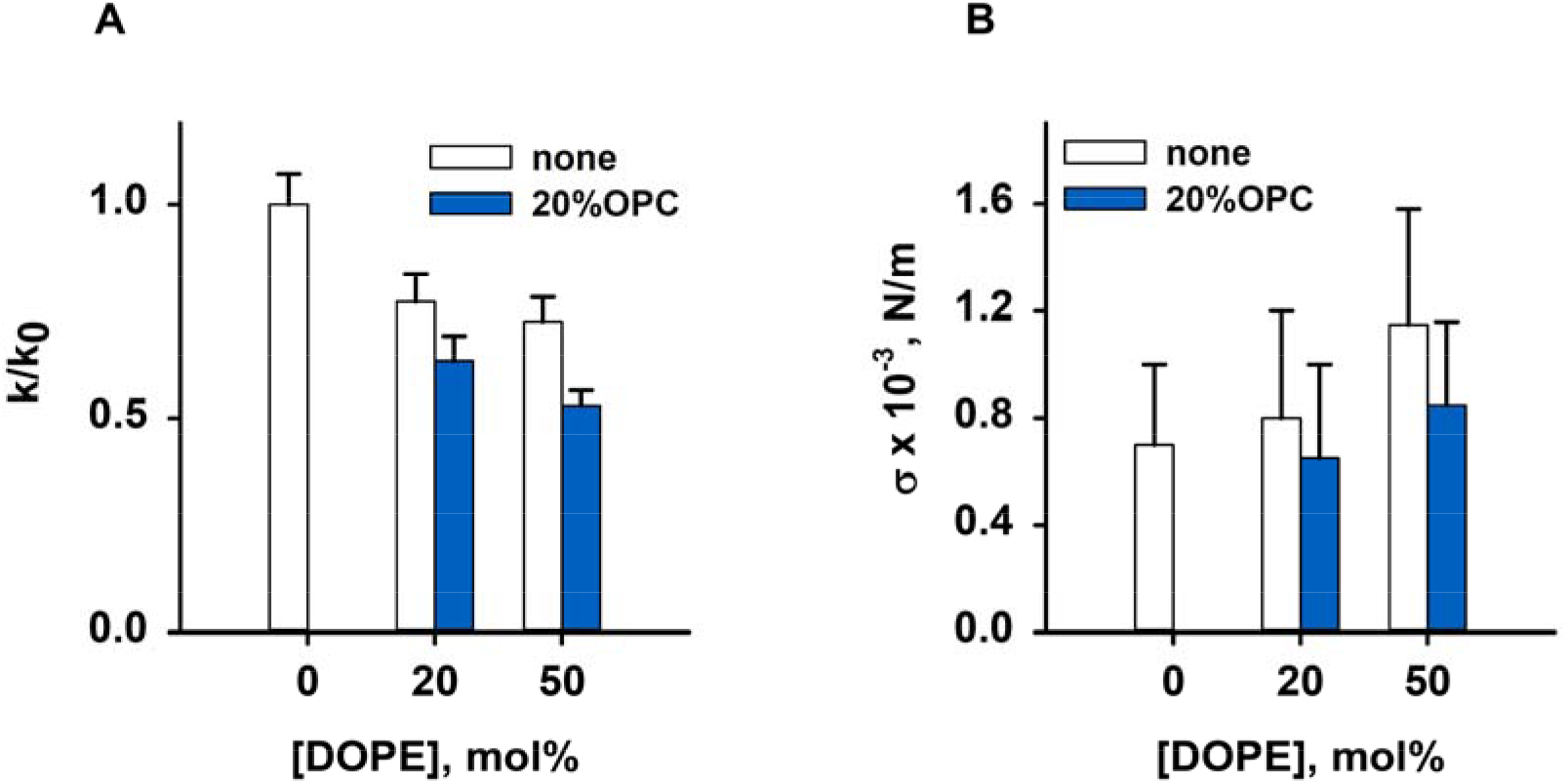
Impact of OPC on the elastic parameters *k/k*_*0*_ and *σ*. C. The lipids were added simultaneously in the ratio as indicated in the diagrams. Buffer composition was as described in Figure 2. Data points represent mean and standard deviation from more than 10 independent experiments.

Alongside enhanced compliance to bending, we observed a decrease in the *σ* for PE-containing membranes after their incubation with RAs (Fig. 2A). Again, this effect was absent in PE-free membranes (Fig. 2B). MD simulations predicted a significant alteration of the PE molecular shape caused by its modification by RAs ^37^. Such lipid reshaping is highly likely to be the reason for the simultaneous reduction in both the *k* and *σ* caused by RAs in PE-containing membranes.

### 3. OPC alters the elastic properties of the lipid bilayer membrane

Finally, we tested the impact of inverted conical lipid 1-hydroxy-2-oleoyl-sn-glycero-3-phosphocholine (OPC; SC > 0) on the elasticity of PE-containing membranes. The presence of OPC mimics the accumulation of lysolipids in the IMM due to increased PLA2 activity in response to oxidative stress. Therefore, we measured the changes in the *σ* and *k* of a membrane comprising the cylindrical lipid DOPC (SC ≈ 0) induced by the addition of DOPE (SC < 0) alone or simultaneously with OPC (SC > 0) in the molar ratios indicated in the figure description (Fig. 3 and Suppl. Fig. 1C and D). Our results showed that the presence of both conical lipids made membranes more compliant to bending than DOPE alone. The bending modulus of a DOPC membrane containing 20 mol% PE and 20 mol% OPC (*k = (13*.*6 ± 0*.*63) kT*) was lower than that of a DOPC:DOPE (1:1) lipid mixture (*k = (16*.*7 ± 1*.*05) kT*). Thus, the OPC-induced changes in elastic parameters in PE-containing membranes were similar to those caused by the accumulation of PE-RAs adducts in the membrane. Both OPC and PE-RA adducts diminished the SCES initially produced by PE.

Moreover, the replacement of DOPC with DOPE led to a concentration-dependent increase in the lateral tension σ in the lipid bilayer membrane (Fig. 3B), as previously shown ^40^. The replacement of 20 mol% DOPC with lysolipid OPC abolished the increase in the σ induced by DOPE. This indicates that the SCES-compensating effect of OPC was probably due to the complementarity of DOPE and OPC shapes (Fig. 3B and Suppl. Fig. 1D).

### 4. The IMM proteins UCP1 and ANT1 sense stored curvature elastic stress

To test our hypothesis concerning the impact of SCES on the function of the mitochondrial membrane proteins, we reconstituted UCP1 or ANT1 in lipid bilayer membranes containing lysolipids with pronounced positive individual curvature, namely, 1-myristoyl-2-hydroxy-sn-glycero-3-phosphocholine (MPC; 14:0) and OPC (18:1). Because the acyl chain length determines the intrinsic lipid curvature (the shorter the acyl chain, the higher the individual lipid curvature), we achieved different SCESs in the lipid bilayer membrane at the same concentration of lysolipids. We compared the specific conductance G_m_ of membranes comprising (i) DOPE:DOPC:CL (45:45:10 mol%), (ii) MPC:DOPC:DOPE:CL (5:45:40:10 mol%), or (iii) OPC:DOPC:DOPE:CL X:(45-X/2):(45-X/2):10, (X=5, 10, and 12.5 mol%), thereby keeping a constant protein to lipid ratio. In all experiments, the UCP1 and ANT1 activity was measured in the presence of the arachidonic acid (AA; 20:4, ω6). With 5 mol% MPC, the G_m_ (MPC) increased to 124.6 ± 13.7 nS/cm^2^, in contrast to the G_m_ = 79.9 ± 5.3 nS/cm^2^ in the absence of MPC. In the experiments with OPC, we demonstrated a concentration-dependent increase in G_m_, from G_m_ =89.9 ± 6.7 nS/cm^2^ in the absence of OPC to the G_m_(OPC) = 158.5 ± 15.8 nS/cm^2^ at 12.5 mol% OPC (Fig. 4B). We failed to measure the conductances of the protein-containing membranes at lysolipid concentrations higher than 5 mol% MPC or 12.5 mol% OPC because of the high membrane instability. In the control measurements, we showed that, in the absence of AA, even 15 mol% OPC does not affect UCP1-mediated G_m_, which remained comparable to the G_m_ (OPC) = 11.2 ± 1.5 nS/cm^2^ of neat lipid bilayer membranes (Suppl. Fig. 2A). The relative increase in G_relANT1_ = 1.65 (G_relANT1_ = G_mANT1_/G_m0_), mediated by ANT1 reconstituted in lipid bilayer membranes containing 10 mol% OPC, was comparable to the G_relUCP1_ = 1.5, measured for the membranes reconstituted with UCP1 in the presence of OPC (Fig. 4B and Suppl. Fig. 2).

**Figure 4.**
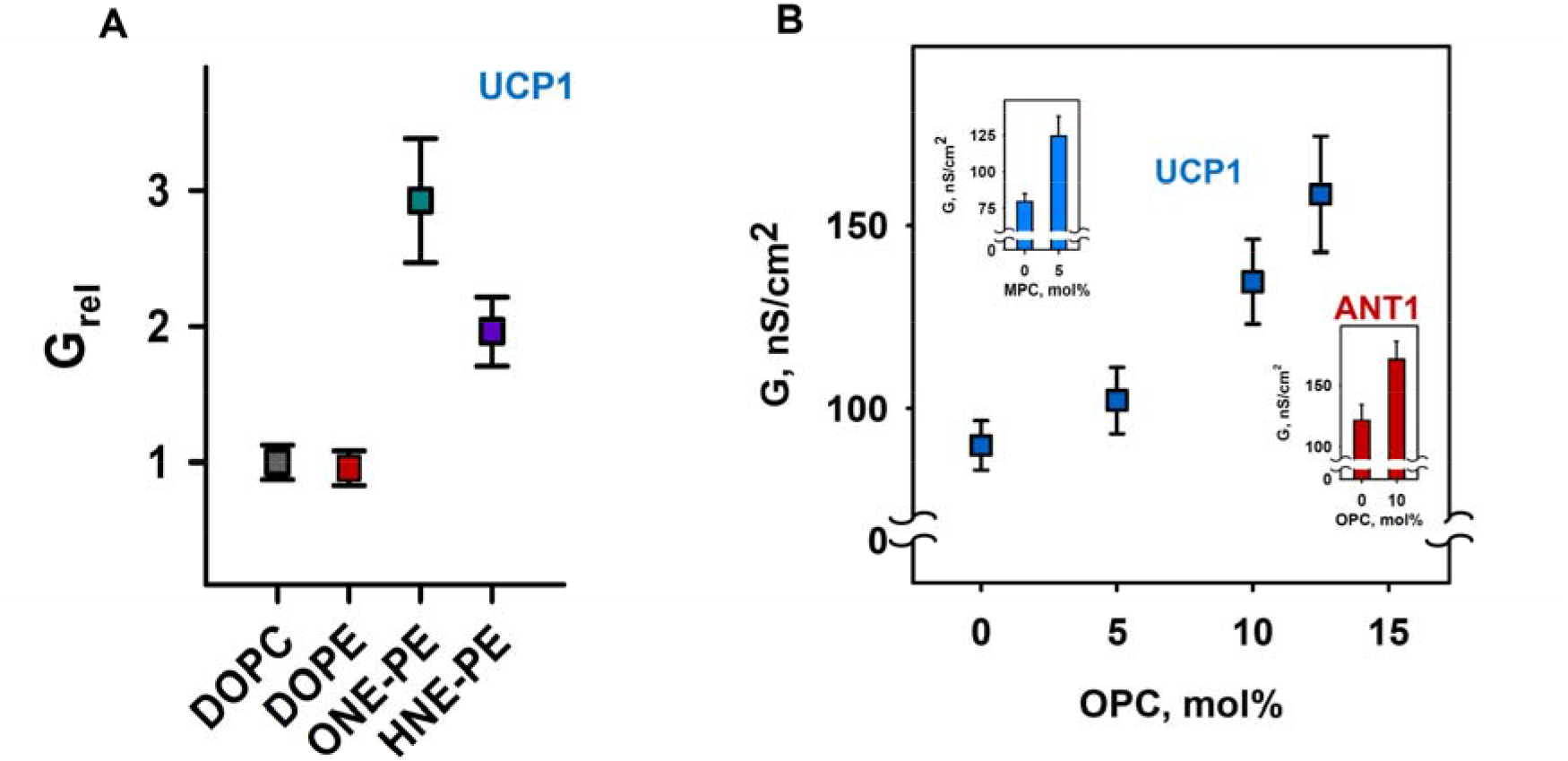
Lipid shape affects UCP1 and ANT-mediated total membrane conductance (*G*_*m*_). A. Influence of PE adducts, ONE-PE (dark cyan), and HNE-PE (violet) on the UCP1-mediated *G*_*m*_ in the presence of AA. DOPC and DOPE were used as controls. Data are taken from Jovanovic et al. ^33^. The relative membrane conductance G_rel_ was calculated as the quotient of the obtained specific membrane conductance in the presence of PE or RA-PEs. B. Increase in the *G*_*m*_, mediated by UCP1 (blue) and ANT (red, insert) in the presence of AA depends on the concentration of the lysolipids OPC and MPC (insert). In control experiments, the lipid composition was DOPC:DOPE:CL 45:45:10. The specified lysolipid amount (mol%) was used instead of DOPC and DOPE. In all experiments, the protein concentrations of UCP1 and ANT1 were 4-5 µg/mg. Concentrations of lipid and AA were 1.5 mg/ml and 15 mol%, respectively. The buffer solution contained 50 mM Na_2_SO_4_, 10 mM MES, 10 mM Tris, and 0.6 mM EGTA, at pH 7.32 and 32 °C. Data points represent means and standard deviation from 3–5 independent experiments.

We tested how SCES affects the inhibition of ANT1 by adding 4 mM ATP to the ANT1 reconstituted with 10% OPC (Suppl. Fig. 2C). A slight decrease in the relative ANT1 inhibition (□ 66% in the presence of OPC vs. □ 76% in its absence; Suppl. Fig. 2C) suggests that the binding of ATP to R79 of ANT1 ^48^ was unaffected by SCES. We were unable to reconstitute the planar lipid membrane with 20 mol% MPC or OPC, even without proteins, which indicates the upper limit of lysolipids (15 mol%) for IMM model lipid composition.

### 5. Impact of lipid shape on the lateral pressure profiles across the lipid bilayer membrane

The results shown above suggest that lipids with different intrinsic curvatures promote activation of the membrane proteins to different extents. Therefore, we used MD simulations to investigate the impact of lipid shape on the LPP in lipid membranes comprising (i) DOPC, (ii) DOPC:ONE-DOPE, (iii) DOPC:OPC, and (iv) DOPC:DOPE (Fig. 5, A-C and Suppl. Fig. 3). The ONE-DOPE adduct is chosen as a prototypical Schiff base adduct formed via the reaction of ONE with PE ^33^. The LPP arises due to the repulsive interaction in the lipid headgroup and acyl chain regions and strong attraction at the lipid-water interface.

**Figure 5.**
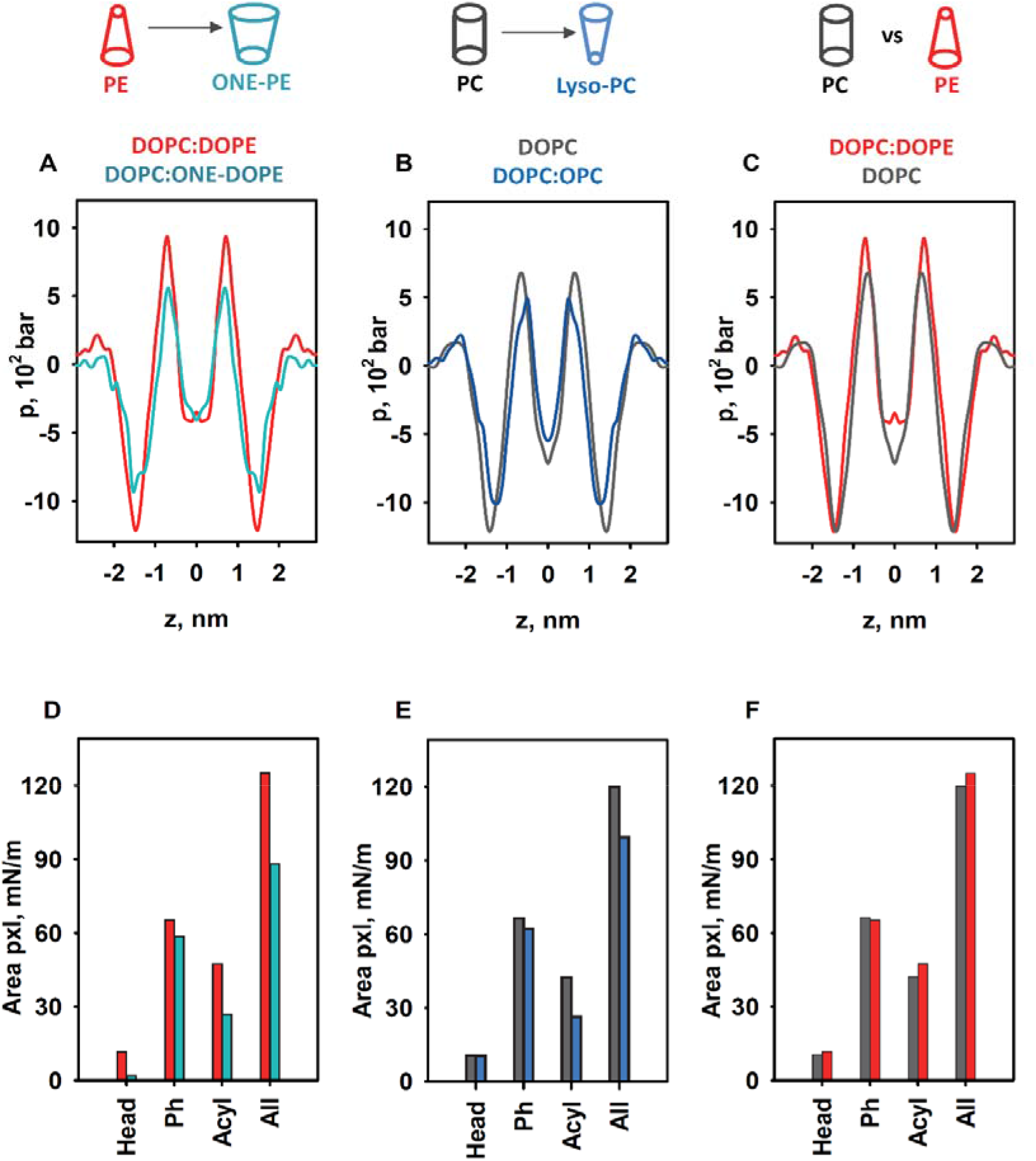
MD simulations of LPP dependency on lipid shape. A-C. Impact of lipids with pronounced SC on the LPP, *p*. D–E. Comparison of areas below the pressure profiles in the headgroup (Head), phosphate group (Ph), and acyl chains (Acyl) and for the whole bilayer profile (All) for *p* shown in A-C. The lipid membrane composition as indicated: (A and C) DOPC:DOPE, (A) DOPC:ONE-DOPE, (B and C) DOPC, and (B) DOPC:OPC. The lipid ratio was 50:50 except for that of the DOPC lipid bilayer membrane. Color labels: DOPC:DOPE (red), DOPC (grey), DOPC:ONE-DOPE (cyan), and DOPC:OPC (blue).

To highlight the fact that the changes originated from the modified PE headgroup due to the formation of the RA-PE adducts, we compared the LPPs in DOPC:DOPE and DOPC:ONE-DOPE membranes (Fig. 5A). A substantial drop in the pressure occurred across the whole membrane profile, in the headgroup region, hydrophobic core, and water-lipid interface. In the membrane-water interface region, Δp_ONE-PE_=(p_PE_ - p_ONE-PE_) was equal to 302 bar (25%), while in the acyl chain region, the Δp_ONE-PE_ was 72.5 bar (40%). To evaluate the contribution of lysolipids to the LPP, we compared the DOPC and DOPC:OPC membrane bilayers (Fig. 5B). The decrease in lateral pressure due to the replacement of DOPC with OPC occurred over the phosphate and acyl tails regions and was slightly pushed toward the center of the bilayer. Compared to the LPP of pure DOPC membranes, the lateral pressure in the OPC-containing membranes (Fig. 5B) decreased in the membrane-water interface region, from Δp_OPC_ = (p_DOPC_-p_OPC_) = 188.6 bar (15.76%), to Δp_OPC_ = 182 bar (27%) in the acyl chain region, and to *Δ*p_OPC_ = 38.6 bar in the middle of the bilayer. We also analyzed the impact of the DOPE (SC < 0) on the LPP in the lipid bilayer by comparing the DOPC:DOPE membrane bilayer with that of DOPC alone, with almost neutral curvature (SC ≈ 0). In contrast to OPC and ONE-PE, insertion of DOPE into the lipid bilayer slightly increased the lateral pressure in the acyl chain region (Δp_DOPE_ = (p_DOPE_ - p_DOPC_) = 256.2 bar [27.5%]), and slightly shifted the profile outwards from the bilayer center (Fig. 5C). Importantly, the alterations in the LPP caused by the insertion of DOPE, ONE-PE, and OPC into the DOPC bilayer are entirely in agreement with the lateral tension measurements (Figs. 2 and 3).

We quantified the distribution of the LPP for each lipid bilayer by calculating the area under the pressure profile curve for the headgroup, phosphate, and hydrophobic regions and compared unmodified lipids to modified lipids (PE vs. ONE-PE, DOPC vs. OPC, and PE vs. PC) (Figs. 5D –F).

Our results showed that the transformation of the PE headgroup (PE_→_ONE-PE) significantly decreased the area below the curve (29.6%), while the deletion of the acyl chain (DOPC_→_OPC) reduced the calculated area by 17% (Figs. 5D and 5E). More precisely, for the PE_→_ONE-PE and DOPC_→_OPC modifications, the drops in the area below the curve were 10% and 6.5% in the phosphate group region and 43.1% and 37.6% in the acyl chains region, respectively (Figs. 5D and 5E). In contrast, PE insertion into the lipid bilayer slightly increased the area below the lateral pressure curve by 4.3% in total, and by 11.9% specifically in the acyl chain (Fig. 5F). These observations are in the agreement with the data shown in Zoni et al. ^49^ that were obtained for LPPs and areas below the curve in the hydrophobic core region for mixed bilayers containing DOPC, DAG, DOPE, and DLPC, where DLPC can be considered a lipid with a positive individual curvature ^17^. Based on the above analysis, the individual curvature of the ONE-PE should be more like that of OPC and, in any case, substantially different from the shape of the endogenous PE. Taken together, our results suggest that membrane protein activity increases if the lipid modification decreases the lateral pressure in the phosphate and acyl chain regions, as shown in neat lipid bilayers; otherwise, it stays unchanged.

## Discussion

The importance of the membrane lipid shape/reshaping for the mitochondria functioning under oxidative stress conditions can be summarized in two related categories: (i) softening of the lipid bilayer membrane, and (ii) a regulatory role for membrane proteins.

Here, we show that the integral proteins, UCP1 and ANT1, embedded in IMM-like membranes, sense changes in SCES caused by oxidative stress. The experimental results and MD simulations revealed that the modified phospholipids, RA-PE adducts, and lysolipids induced similar changes in the bending rigidity, lateral tension, and LPP of PE-containing membranes. The observed variations in *k* and *σ* among RAs can be explained by differences in the types of RA-PE adducts and their localization in the lipid bilayer membrane ^33^. Because OPC (SC > 0) induced similar changes in *k, σ*, and p as a ONE-PE adduct in PE-containing membranes, we speculate that this PE adduct has a positive individual curvature. Thus, RA-driven shape transformation in a fraction of PE molecules with a negative SC reduced lipid packing stress due to mutual compensation with lipids with the opposite SC, thereby decreasing the membrane SCES.

A zwitterionic lysoPC lipid increased UCP1- and ANT1-mediated proton translocation similarly to RA-PEs. More importantly, the impact of RA-PEs on the elastic parameters k and σ in the order HHE<HNE<ONE perfectly matches with their ability to potentiate UCP1-mediated proton translocation in the presence of FFAs in the membranes of the same lipid composition (Fig. 4A, right) ^33^. This supports the hypothesis that RA-induced changes in a lipid environment, and not a modification of protein amino acid residues, increases the G_m_. Thus, we propose that lipid shape and, with it, related membrane mechanical properties, such as bending rigidity and lateral pressure, play a regulatory role in the protonophoric activity of UCPs and ANTs. In contrast, membrane surface potential, which increases due to the formation of RA-PEs ^33^, is most probably irrelevant for UCP1 and ANT1 potentiation because zwitterionic lysolipids do not affect the surface potential.

Notably, MD simulations suggested decreased lateral pressure over the whole membrane profile for lipid bilayer membranes composed of ONE-PE adduct or OPC. We hypothesized that, at the molecular level, the altered lipid environment results in increased G_m_ mediated by UCP1. Lipid environment changes are caused by the type of lipid transformation in which the ratio (lipid head)/(acyl chains) is increased compared to the lipid “initial form” (e.g., DOPC_→_OPC, PE_→_RA-PE). These modifications decreased both the bending modulus *k* and lateral pressure *p* in the lipid bilayer membrane (Figs. 2A, 5A and 5B).

Although the insertion of PE into the PC lipid bilayer induced a significant decrease in the bending modulus k (Fig. 1D, 2, and Suppl. Fig. 1A and C), it slightly increased the LPP in the acyl chain region (Fig. 5C and F). This resulted in a lateral tension increase (Fig. 4), whereas the UCP1-mediated conductance stayed unchanged (Fig. 1A, left). In the case of RA-PE and lysolipids, a decreased lateral pressure at the acyl tail region might potentiate the probability that anionic FFAs will (i) reach the protein binding site, which is localized to the hydrophobic region close to the lipid phosphate groups on the matrix side of the IMM ^48^, (ii) be protonated in the position close to the bilayer center (asterisk in Fig. 6C and ^50^), and leave protein. Eventually, the same lipid environment supports the protein conformation change, ensuring faster FFA^-^ translocation. Notably, protein modifications, such as crosslinking of RA-amino acid adducts or protein mutations, can lead to loss of the protonophoric function, whereas lipid modification potentiates the protein-mediated FFA^−^ translocation. Remarkably, in the absence of the oxidative stress lipids with distinctly positive curvature, such as lysolipids and PIPs, are found only in trace amounts (< 1%) and are primarily involved in direct lipid–protein interaction and signaling ^6^.

**Figure 6.**
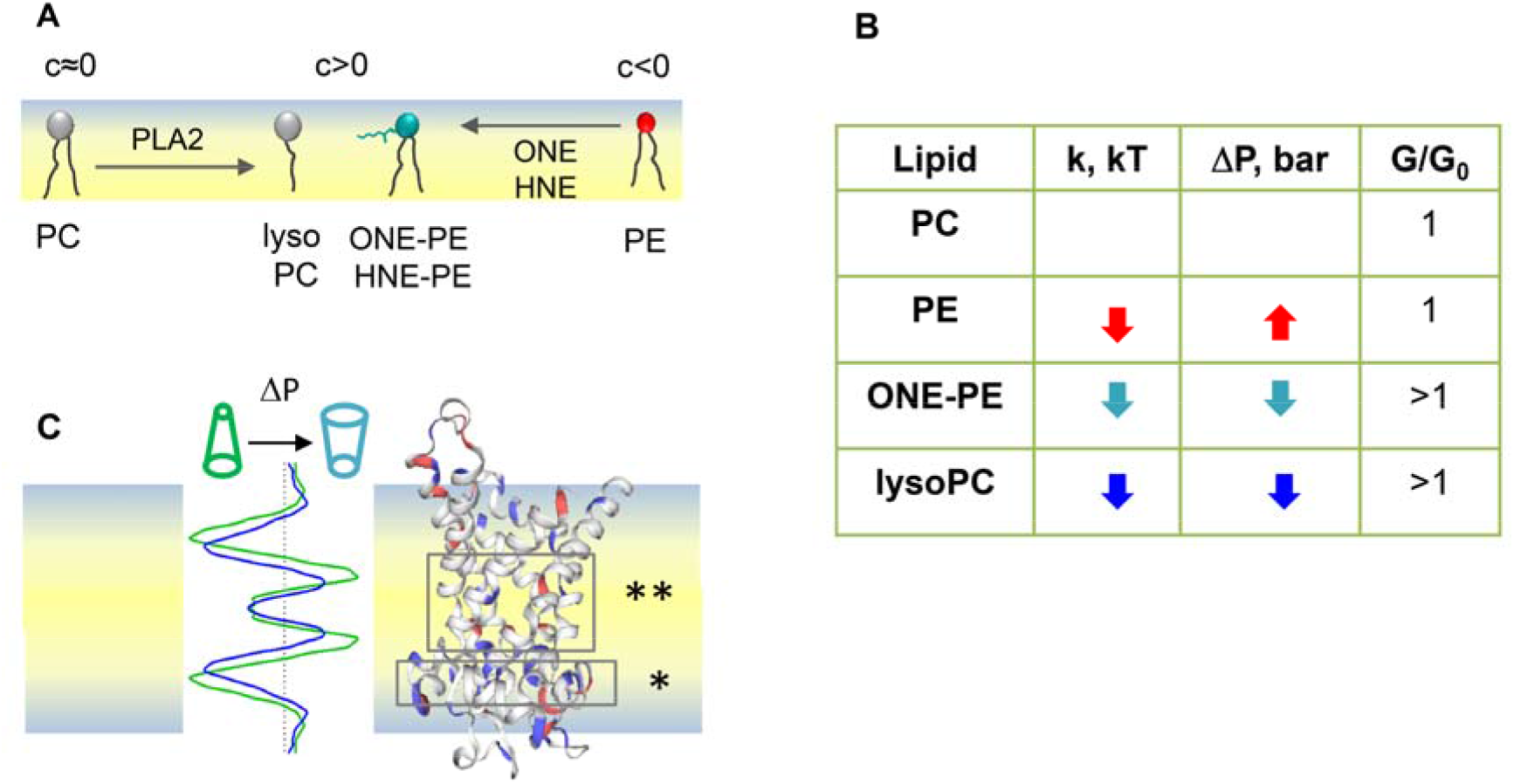
Proposed mechanism for the SCES sensing by mitochondrial proteins ANT1 and UCP1. **A**. Schematic representation of the lipid membrane leaflet and processes that cause a change in their shape during oxidative stress in mitochondria. **B**. The increase in the UCP1- and ANT1-mediated G_m_ correlates with a decrease in both, the lateral pressure P and bending rigidity k in the lipid bilayer membrane. If the change in P and k goes in the opposite direction, the protein-mediated G_m_ is not affected, as demonstrated here for the DOPE-containing membrane. **C**. Schematic representation of the lipid bilayer membrane and distribution of the LPP caused by a change in the lipid shape. A comparison of the LPP (left) and protein structure (right) suggests that decreased lateral pressure (cyan), which appeared in the area of the FFA binding site (*) and in the protein cavity region (**), promotes FFA-translocation. Most likely, the transformed lipid environment (i) improves the probability that anionic FFA will reach a protein binding site and release protein and (ii) facilitate a protein conformational change, which in turn supports a faster translocation of anionic FFA to the opposite leaflet.

## Conclusion

Identification of the mechanisms by which the transformation of lipid shape affects the functioning of mitochondria helps to explain the onset of diseases associated with oxidative stress in a way that has not been considered so far. Our results show that lysolipids and PEs modified by RAs similarly affect membrane mechanical properties by decreasing the bending modulus *k* and SCES. Furthermore, we showed that UCP1 and ANT1 sense SCES and proposed a new mechanism for regulating the protonophoric function of IMM proteins under oxidative stress conditions.

## Materials and methods

### 1. Chemicals

1,2-dioleoyl-sn-glycero-3-phosphocholine (DOPC), 1,2-dioleoyl-sn-glycero-3-phosphoethanolamine (DOPE), 1-oleoyl-2-hydroxy-sn-glycero-3-phosphocholine (OPC), bovine heart cardiolipin (CL), arachidonic acid (AA), hexane, hexadecane, KCl, HEPES, EDTA, Na_2_SO_4_, MES, Tris, adenosine 5-triphosphate (ATP), and guanosine 5-triphosphate (GTP) were purchased from Sigma-Aldrich (Munich, Germany). MPC (1-myristoyl-2-hydroxy-sn-glycero-3-phosphocholine) was obtained from Avanti Polar Lipids (Alabaster, AL). Plain silica beads (22-μm diameter) were from Microspheres-Nanospheres (Corpuscular Co., Cold Spring, NY). Chloroform was purchased from Carl Roth (Karlsruhe, Germany). 4-oxo-trans-2-nonenal (4-ONE) and 4-hydroxy-trans-2-nonenal (4-HNE) were purchased from Cayman Chemical (Ann Arbor, MI, USA) or synthesized as previously described ^51^.

### 2. Reconstitution of UCP1 and ANT1 in liposomes

Recombinant uncoupling protein (murine UCP1) and adenine nucleotide translocase 1 (murine ANT1) were purified from *E. coli* inclusion bodies and reconstituted into liposomes as previously described ^52, 53^. AA (20:4, ω6), at a concentration of 15 mol%, and lyso-PCs (OPC and MPC), at the concentrations indicated in the figure descriptions, were added to the lipid phase before membrane formation.

### 3. Formation of the planar lipid bilayer membranes and membrane electrical parameters measurements

Planar lipid bilayers were formed from proteoliposomes on the tip of a one-way plastic pipette ^54^. The dispensable container was filled with 0.75 ml buffer containing 50 mM Na_2_SO_4_, 10 mM MES, 10 mM Tris, and 0.6 mM EGTA at pH 7.32 and 32°C. Membrane formation and bilayer quality were monitored by capacitance measurements. Current–voltage (I–V) characteristics were measured by a patch-clamp amplifier (EPC 10; HEKA Elektronik Dr. Schulze GmbH, Germany). Total membrane conductance *G*_*m*_ was calculated from a linear fit of experimental data (I) at applied voltages (*V*) in the range of −50 mV to 50 mV ^55^. The relative conductance, G_rel_, was calculated according to equation Eq. 3:

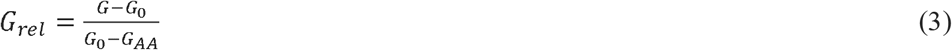

where G and G_0_ are the total membrane conductances of the lipid membranes reconstituted with transmembrane protein and AA, with or without lysolipids, respectively, while is G_AA_ is the total membrane conductance of the lipid membranes reconstituted with AA alone.

### 4. Measurements of the membrane elastic parameters

BLMs comprised DOPC, DOPE, CL, and OPC in the indicated ratios. The BLMs were formed by applying the “painting” technique to the apertures of a grid mesh fixed tightly at a small distance from the bottom of a Petri dish ^45^. The experimental chamber was filled with buffer containing 100 mM KCl, 10 mM HEPES, and 1 mM EDTA at pH 7.0 and room temperature (RT). The apertures of the mesh were pre-treated with a small drop of the lipid mixture (10 mg/ml total lipid) dissolved in decane:octane (1:1 v/v). The solvents were evaporated under an argon stream. Afterward, a small amount of lipid mixture dissolved in squalane (20–30 mg/ml, total lipid) was deposited (“painted over”) on the mesh apertures. “Overpainted” BLMs are formed spontaneously from a thick film covering the grid apertures. The excess lipid material was forced out to the periphery, forming a “toroidal meniscus” – a reservoir, maintaining the lateral tension of the lipid bilayer (Fig. 1A). Nanotubes (NTs) were pulled from BLMs vertically using a fire-polished borosilicate patch pipette with a tip diameter of ∼1 μm filled with the same buffer as the experimental chamber. The tip of the pipette was put into tight contact with the BLM. The hydrostatic pressure pulse ruptured the small membrane patch isolated inside the pipette while the pipette rim remained in contact with the parent BLM. The NT formed spontaneously when the pipette was slowly moved away from the membrane. A precise nanopositioning system (piezo linear actuator and Actuator Controller ESA-CSA, Newport) controlled the vertical position of the pipette (ΔL). The formation of the NT was detected by conductance measurements using Ag/AgCl electrodes placed in the pipette and bath solution (see below). The quality of the NT bilayer membrane was controlled by capacity measurements, whose characteristic value was about *c*_*sp*_ ≈ 1 μF·cm^-2 56^. The NT radius was recalculated from the ion current *I*, measured by an Axopatch 200B amplifier (Molecular Devices, Sunnyvale, CA), and acquired by a low-noise data acquisition system (Axon Digidata 1550, Molecular Devices, Sunnyvale, CA). The amplifier was set in the voltage-clamp mode (fixed voltage) to measure the ion conductance *G* = *I/U*. We used the hyperbolic approximation of G(ΔL) to fit the measured conductance dependence described in ^45^. Briefly, the conductance of the NT lumen, *G*_*NT*_, was obtained as the difference between the *G*_*m*_ and the patch conductance, *G*_*p*_ (*G*_*p*_ = *G* at infinite NT length): *G*_*NT*_ = *G* – *G*_*p*_. The vertical asymptotes of the fit provided the length measurement offset, which should be subtracted from *ΔL* to measure the length of NT (*L*_*NT*_). NT radius (*r*_*NT*_) was determined as given in Eq. 4,

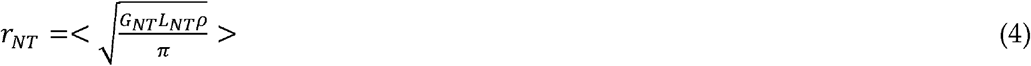

where the *ρ* denotes the specific electrical resistance of the electrolyte we used.

### 5. Molecular dynamics simulations

We performed molecular dynamics (MD) simulations for lipid bilayers of different compositions – DOPC, DOPC:DOPE 50:50, DOPC:ONE-PE 50:50, and DOPC:OPC 50:50. DOPC, DOPE, ONE-PE Schiff base adduct ^33, 37^, and OPC were described with Slipids Force Field ^57, 58, 59^. All missing bonding and non-bonding parameters of lipid molecules in the existing Slipids Force Field were updated with compatible CHARMM36 parameters ^60^ when needed. Atomic charges were recalculated by the standard Slipids procedure using the Merz-Singh-Kollman scheme ^61^, which is composed of B3LYP/6-31G (d) geometry optimization of the molecule of interest, a subsequent single point ESP charge calculation using the B3LYP/cc-pVTZ method, and a final charge refinement with the RESP method ^62^.

Bilayers containing 128 lipid molecules were constructed from two monolayers containing 64 individual lipid molecules, with symmetrical lipid distribution across the leaflets in mixed systems. All systems were placed in a unit cell and solvated by about 12,000 water molecules using the TIP3P water model ^63^. The unit cell size was approximately 6.5 × 6.5 × 12.0 nm. 3D periodic boundary conditions were used with long-range electrostatic interactions beyond the non-bonded cut-off of 1 nm using the particle-mesh Ewald procedure ^64^ with a Fourier spacing of 1.2 nm. Real-space Coulomb interactions were cut off at 1 nm, while van der Waals interactions were cut-off at 1.4 nm. We performed 100-ns MD simulations with semi-isotropic pressure coupling, independently in the directions parallel and perpendicular to the bilayer’s normal using the Parrinello–Rahman algorithm ^65^. The pressure was set to 1 bar, and a coupling constant of 10 ps^-1^ was used. All simulations were performed at 310 K and controlled with the Nose–Hoover thermostat ^66^ independently for the lipid-water sub-systems, with a coupling constant of 0.5 ps^-1^. Bond lengths within the simulated molecules were constrained using the LINCS ^67^. Water bond lengths were kept constant by using the SETTLE method ^68^. Equations of motion were integrated using the leap-frog algorithm with a time step of 2 fs. For the LPP, a custom version of GROMACS-LS package ^69, 70^ was used to re-run trajectories and output local stress tensors. Because long-range electrostatics via PME is not available in GROMACS-LS, an increased cut-off distance of 2 nm was used for Coulomb interactions calculations as suggested by the package developers ^69^. Error bars for pressure profiles of lipids were calculated as the difference between symmetrized and unsymmetrized pressure profiles in different leaflets. Smoothed data were obtained from the average of two points.

### 6. Statistics

Data from the electrophysiological measurements are displayed as mean ± standard deviation of at least three technical replicates (on three different days). Each replicate was the mean membrane conductance from a minimum of three formed bilayer membranes.

## Data availability

The data supporting the findings are available in the Supplementary Information and from the corresponding author upon reasonable request.

## Acknowledgments

We are grateful to Sarah Bardakji for the excellent technical assistance and for helping with the production and evaluation of recombinant proteins. We thank the computer cluster Isabella based in SRCE at the University of Zagreb, University Computing Centre for computational resources.

## Authors contributions

O.J and E.E.P. designed the study; O.J, K.C., P.B., and K.Z. performed the electrophysiological experiments; S.Š. and M.V. performed the molecular dynamics simulations; O.J., K.C., P.B., K.Z., and M.V. analyzed the data and prepared the figures; O.J., P.B., and E.E.P. interpreted the data and wrote the manuscript draft. All authors revised the manuscript and agreed on the final version.

