## Supplementary Figures for "Membrane lipid reshaping underlies oxidative stress sensing by the mitochondrial proteins UCP1 and ANT1"

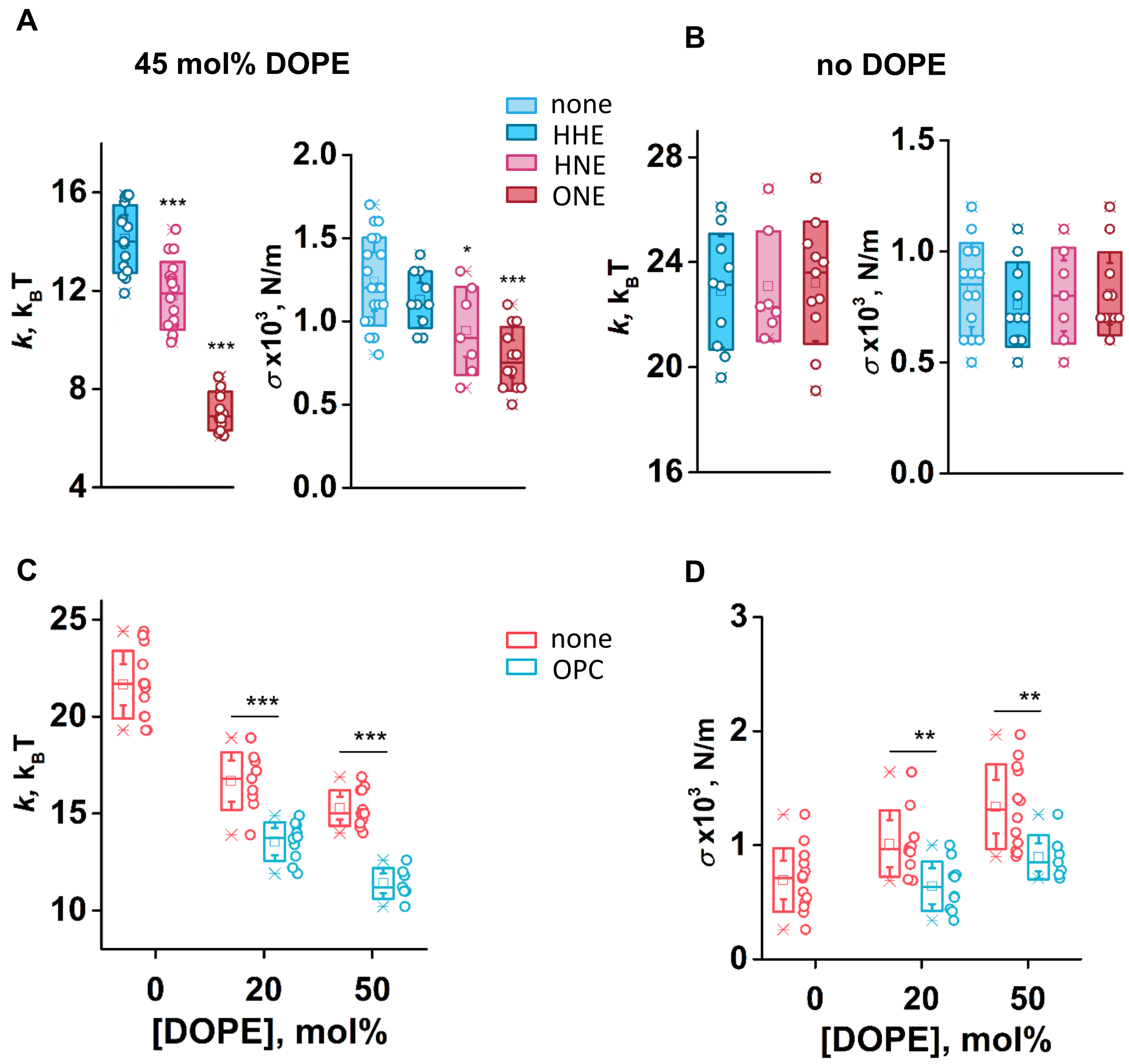


**Supplementary figure 1. Lipid shape alters the membrane's elastic properties.** A and B. Bending modulus k and lateral tension σ measured for lipid membranes composed with (A) and without DOPE (B), incubated with reactive aldehydes (RAs) 4-hydroxy-2-hexenal (HHE), 4-hydroxy-2-nonenal (HNE), and 4-oxo-2-nonenal (ONE). Membrane and buffer composition as described in Figure 2. RAs concentration is in the range of (500 – 700) µM. Changes induced by RAs are significant (* P<0.05; *** P<0,001 t-tests). C and D. The impact of DOPE on k and σ when inserted in DOPC lipid bilayer composed with (pink) and without (blue) OPC. Data are significantly different (** P<0.01; *** P<0.001). Boxes in box-charts indicates mean ± SD, whickers represent mean ± 95%.

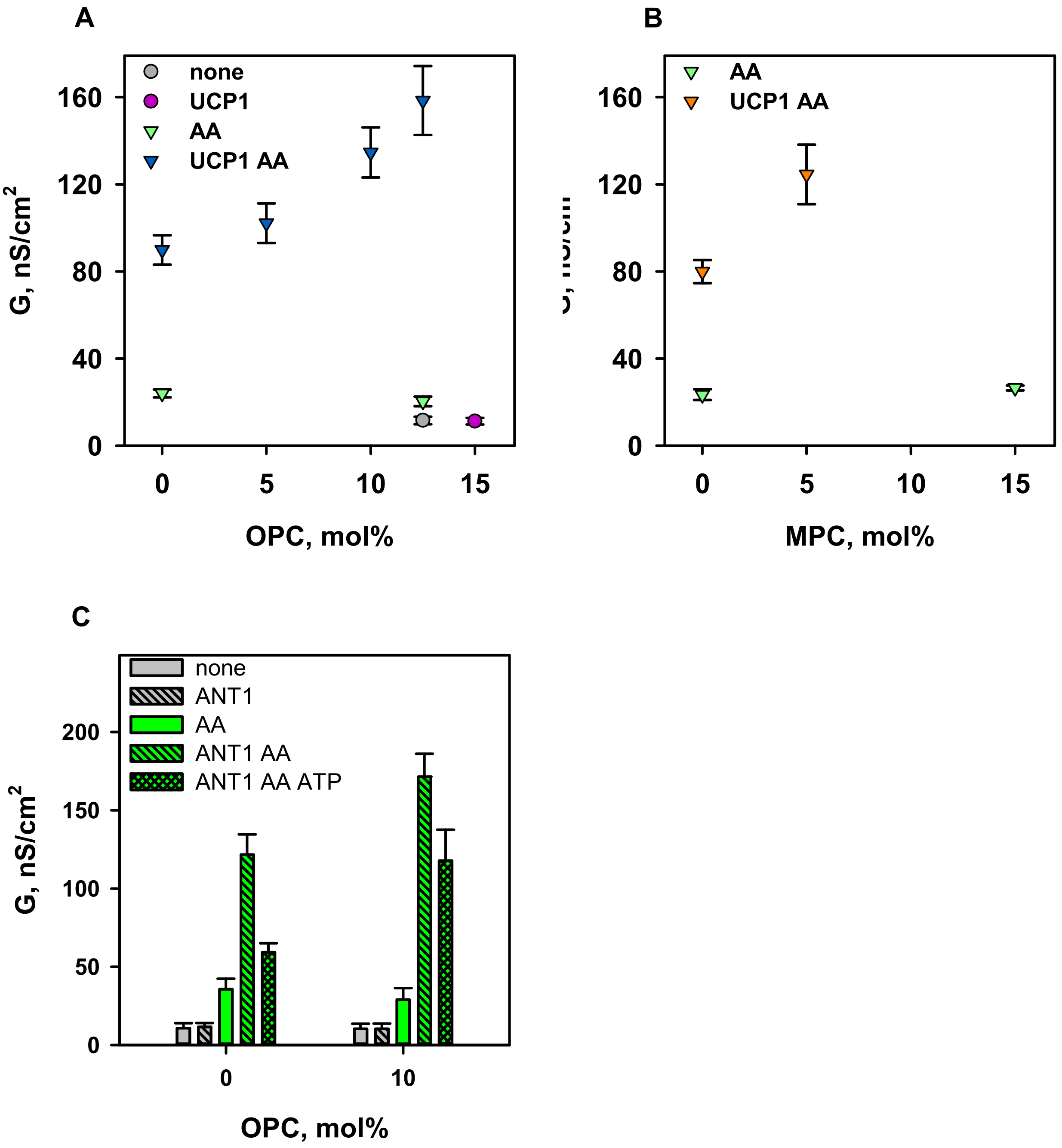


**Supplementary figure 2.** A. Influence of OPC, (18:1)PC, on total membrane conductance G membranes reconstituted with UCP1 (pink) and without (grey), as well with AA, without UCP1, (green) and with UCP1 (blue) respectively. B. Influence of MPC, (14:0)PC, on G lipid bilayer membrane reconstituted with AA in the presence (orange) and absence (green) of UCP1. C. Influence of 10 mol% OPC on G planar lipid bilayer membrane reconstituted: (i) with (grey diagonal pattern) and without (grey) UCP1, (ii) with AA only (green) and with ANT1 (green diagonal pattern), (iii) with ANT1 and AA inhibited by 4mM ATP (green romb pattern). In all experiments, concentrations of lipid and AA were 1.5 mg/ml and 15 mol% respectively. UCP1 concentration was 4-5 µg/mg ANT1 concentration was 4µg/mg. In control experiments, lipid composition was DOPC:DOPE:CL 45:45:10. OPC mol% was taken instead DOPC and DOPE. Buffer solution contained 50mM $Na_2SO_4$, 10 mM MES, 10mM TRIS, 0,6mM EGTA, at pH 7,32 and T=32°C. Data points represent means and standard deviation from 3-5 independent experiments.

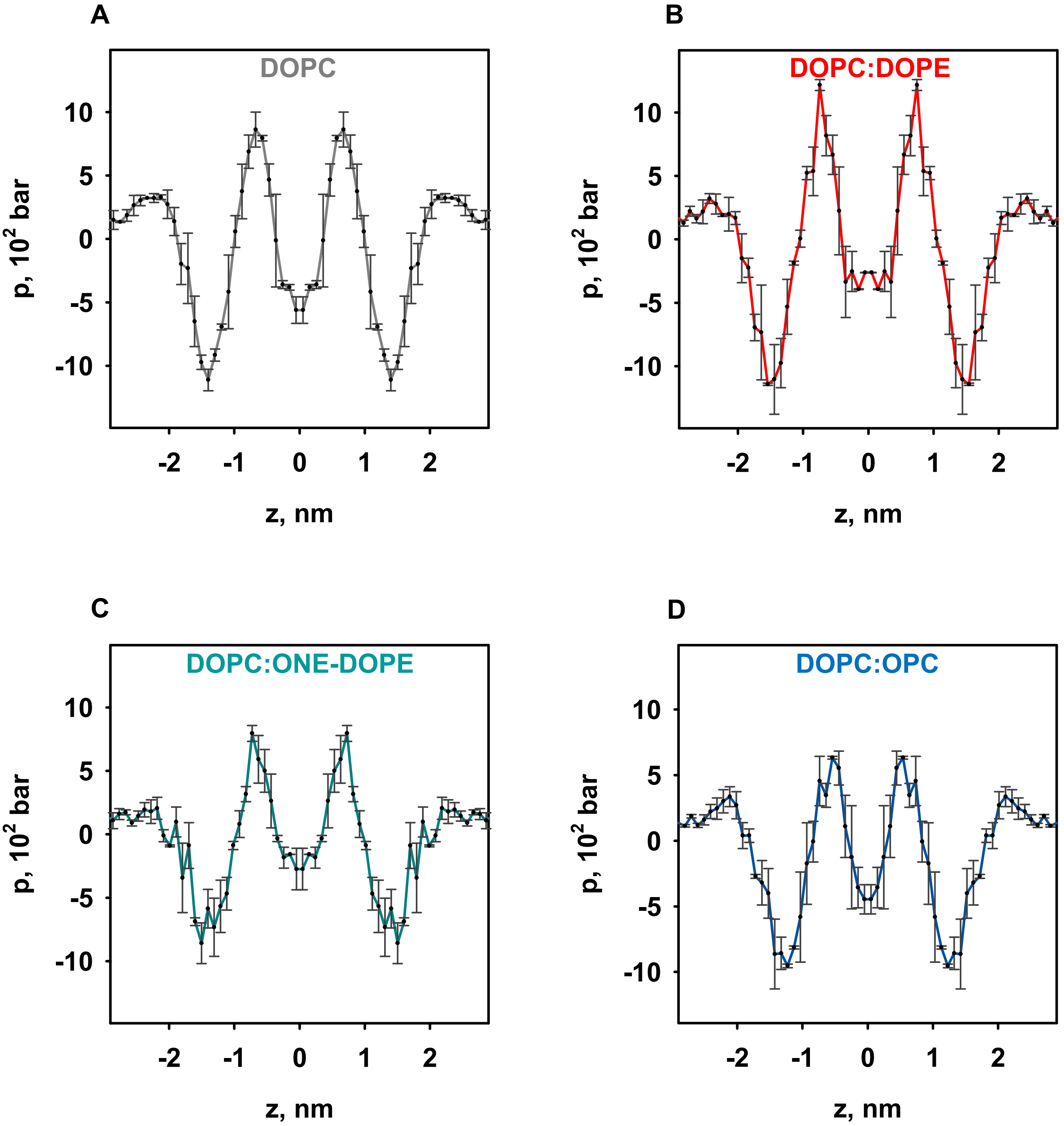


**Supplementary figure 3.** Lateral pressure profiles of DOPC (A), DOPC:DOPE (B), DOPC:ONE-DOPE (C) and DOPC:OPC (D). Error bars were calculated as a difference between symmetrized and unsymmetrized pressure profiles in different leaflets.
